# Layer-specific Loss and Compensation of Parvocellular Response in Subcortical Pathways of Adult Human Amblyopia

**DOI:** 10.1101/2020.04.02.022129

**Authors:** Wen Wen, Yue Wang, Sheng He, Hong Liu, Chen Zhao, Peng Zhang

## Abstract

Abnormal visual experience in critical period causes amblyopia or lazy eye, reducing visual abilities even with corrected optics. A long-standing question is where in the human visual system does the amblyopic deficit arise. In particular, whether amblyopia induces selective deficits of the magnocellular (M) or the parvocellular (P) geniculostriate pathways, and whether the more ancient retinotectal pathway is also affected. Technical limitations to non-invasively measure layer-specific activity in human lateral geniculate nucleus (LGN) and superior colliculus (SC) hampered efforts in addressing these questions. In the current study, using lamina-resolved 3T and 7T fMRI and visual stimuli selectively activating the M and P pathways, we investigated layer-specific response properties of the LGN and the SC of amblyopia patients and normal controls. With stimuli presented to the amblyopic eye, there was a stronger response loss in the P layers than in the M layers of the LGN. Compared to normal controls, amblyopic eye’s response to the P stimulus was selectively reduced in the superficial SC, while the fellow eye’s response was robustly increased in the deep SC. Selective P response deficits of amblyopia were also observed in the visual pulvinar, early visual cortex, and ventral but not dorsal visual streams. These results provide strong in vivo evidence in adult amblyopic patients for selective deficits of parvocellular functions in the visual thalamus, and additionally reveal response deficits to the amblyopic eye and neural compensation to the fellow eye in the retinotectal pathway.

**Highlights:**

- Parvocellular response loss in the LGN P layers, visual pulvinar and ventral visual stream
- Selective amblyopic deficits of the parvocellular pathway
- Amblyopic eye’s response decreased in the superficial SC
- Fellow eye’s response increased in the deep SC
- Amblyopic deficits and neural compensation in the retinotectal pathway

## Introduction

Abnormal visual experience in the critical period, commonly due to a turned eye (strabismus) or unequal refractive errors (anisometropia), causes amblyopia or lazy eye with impaired visual abilities even with corrected retinal input. A long-standing question is where in the human visual system does the amblyopic deficit arise. The current consensus is that amblyopic damage was post-retinal in the subcortical and cortical visual areas of the brain (Hess and Baker 1984, Levi 2006). Human neuroimaging studies found functional deficits of amblyopia in the visual cortex (Anderson and Swettenham 2006, Joly and Franko 2014). However, due to poor neuroimaging signals in the human subcortex, little is known about the amblyopic deficit in the human subcortical pathways.

In primate, retinal information reaches visual cortex mainly through the geniculostriate pathway. The lateral geniculate nucleus (LGN) is mainly composed of six layers of neurons with distinct cell types and functions. Parvocellular (P) cells in the four dorsal layers have smaller cell body, sustained discharge pattern, smaller receptive field, low sensitivity to luminance contrast, and center-surround color opponency; magnocellular (M) cells in the two ventral layers have larger cell body, transient neural response, larger receptive field, and high sensitivity to luminance contrast and motion (Derrington, Krauskopf et al. 1984, Derrington and Lennie 1984, Hubel and Livingstone 1990). Thus the M and P stream-specific visual processing are clearly segregated in the LGN. Without direct access of layer-specific response of the LGN, previous studies showed mixed results for selective dysfunctions of either M or P pathways in amblyopia (Levi and Harwerth 1977, Barnes, Hess et al. 2001, Zele, Pokorny et al. 2007, Hess, Li et al. 2009).

Retinotectal pathway is an alternative and more primitive visual pathway. In primate, about 10% of retinal ganglion cells project to the superficial layers of the superior colliculus (SC) (Perry and Cowey 1984), which sends projections to the Koniocellular layers of the LGN and the inferior (visual) pulvinar (May 2006). The superficial SC also receives direct cortical input from the early visual cortex (V1-V3) and area MT (Fries 1984, Lock, Baizer et al. 2003), mainly processes visual sensory information. Deeper layers of the SC (mainly the intermediate layers) receive multisensory input from the brain stem (Jay and Sparks 1987, Wiberg, Westman et al. 1987), and afference from the frontoparietal cortex, basal ganglia and extrastriate visual cortex (Beckstead and Frankfurter 1982, Fries 1984). The deep SC mainly contributes to attention and premotor control of eye and head movements. The visual pulvinar that receives input from the SC sends projections to cortical targets in occipital, parietal and temporal lobes (May 2006). It plays important roles in visual perception and attention, and regulates information transmission between cortical areas (Purushothaman, Marion et al. 2012, Saalmann, Pinsk et al. 2012, Zhou, Schafer et al. 2016). To the best of our knowledge, the influence of amblyopia to the retinotectal pathway and the visual pulvinar remains largely unexplored.

Investigating pathway-specific dysfunctions of the human subcortex could reveal the neural cites and mechanisms of amblyopia, which is important for developing new tools for better treatment and prognosis of the disease. To achieve this goal, a non-invasive neuroimaging tool is required to measure pathway or layer-specific activity of the human LGN and SC. Recent advances in high resolution fMRI techniques and experimental approaches enabled us to resolve layer-specific functional responses of the human LGN and SC (Zhang, Wen et al. 2015, Zhang, Zhou et al. 2015, Yu, Zhang et al. 2016, Qian, Zou et al. 2020). In the current study, using high-resolution 3T and 7T BOLD (blood oxygen level dependent) fMRI and carefully designed visual stimuli selectively activating the M and P layers of the LGN (the M and P stimulus), we investigated layer-specific responses of the LGN and the SC of adult amblyopia patients and normal controls. Responses of the visual pulvinar and cortical visual areas were also examined.

Our results clearly demonstrated amblyopic deficits selective to the response of P layers of the LGN. The M layers’ response was also reduced, but was much less compared to response loss in the P layers. Selective P response loss was also observed in the visual pulvinar, early visual cortex (V1-V3) and ventral visual stream (V4/VO/LO), but not in the dorsal visual stream (V3ab/MT). Compared to normal controls, the amblyopia eye’s response to the chromatic P stimulus was selectively reduced in the superficial SC, whereas the fellow eye’s response to the P stimulus was robustly and selectively increased in the deep SC. These findings provide strong functional evidence in humans for selective parvocellular deficits in the visual thalamus and visual cortex of amblyopia, and reveal response deficit to the amblyopic eye and neural compensation to the fellow eye in the retinotectal pathway.

## Results

### 1. Amblyopic eye’s response was selectively reduced in the P layers of the LGN

In the 3T fMRI experiment (2mm isotropic voxels), M and P biased visual stimuli (Figure 1A) differed in spatial frequency, temporal frequency, contrast and chromaticity were used to selectively activate the M and P layers of the LGN. The amblyopic eye (AE) and the fellow eye (FE) of unilateral amblyopia patients were tested in separate sessions. The non-dominant eye of normal controls (NE) was also tested. We found selective loss of amblyopia eye’s response to the chromatic P stimulus in the P layers of the LGN. In order to reduce the influence from confounding signals and to replicate the 3T fMRI results, we performed a 7T fMRI experiment at a higher spatial resolution (1.2mm isotropic). High contrast and high spatial frequency achromatic stimulus (achromatic P stimulus, Figure 2A) was monocularly presented to the amblyopic eye and to the fellow eye of amblyopia patients. The achromatic P stimulus presented to the fellow eye strongly activated the P layers of the LGN. Compared to the fellow eye, the amblyopic eye’s response was significantly reduced in the P layers but not in the M layers of the LGN.

**Figure 1.**
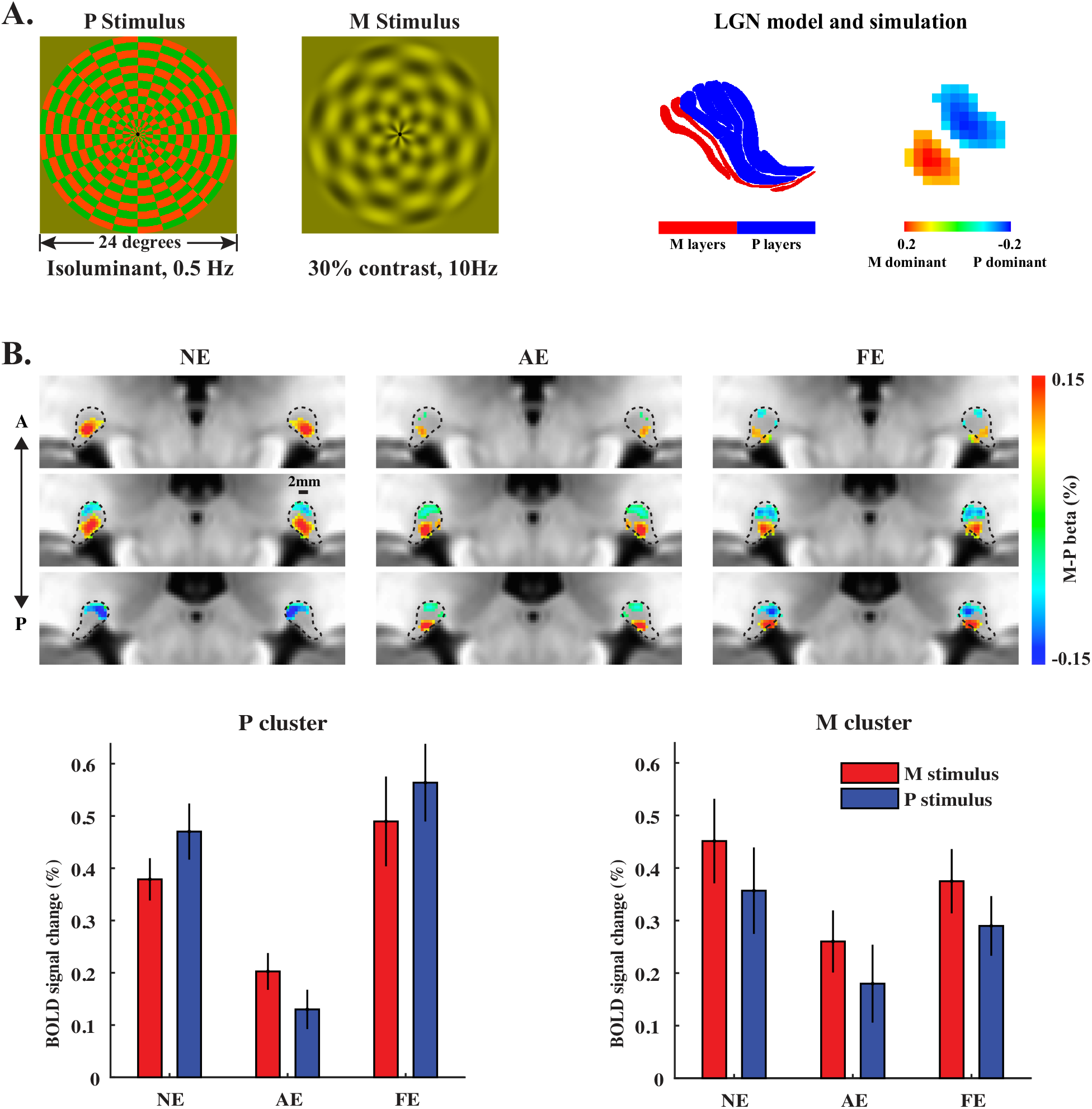
Visual stimuli and results for the 3T fMRI experiment. **A. Left:** M and P biased visual stimuli used in the 3T fMRI experiment. **Right:** Model for the M and P layers of the LGN according to Nissl stained histology of the human LGN (Amunts, Lepage et al. 2013). The right side shows the simulated fMRI pattern for M-P responses. **B. Upper**: M-P beta maps for the M and P dominant clusters. Maps were resampled at 0.6mm isotropic resolution for illustration purpose. Maps for the left and right LGNs are mirror symmetric. **Lower:** BOLD response to the M and P stimuli in the M and P dominant clusters of the LGN. Error bars represent standard error of the mean. Abbreviations, NE: normal eye, AE: Amblyopic eye, FE: Fellow eye.

**Figure 2.**
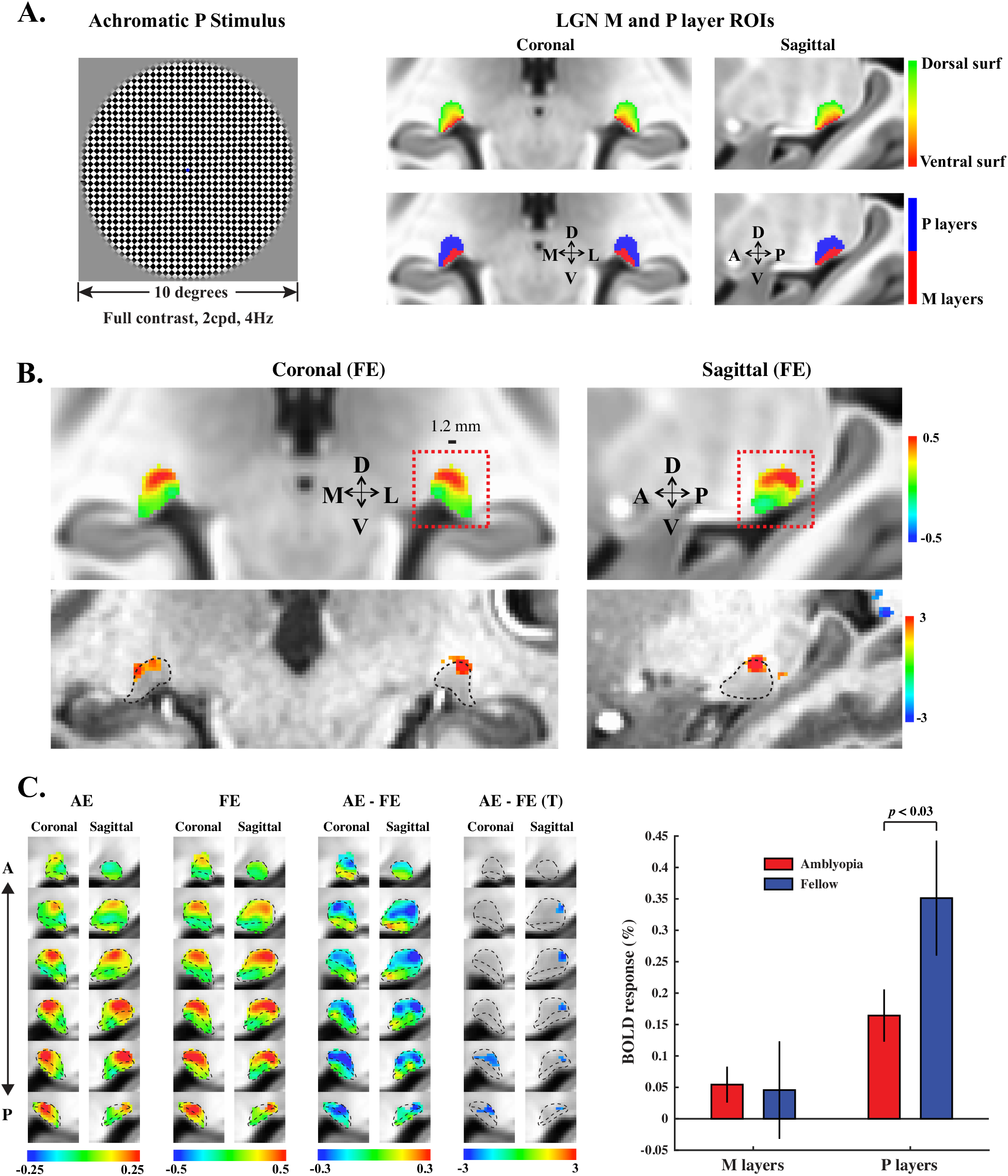
Stimulus and LGN results for the 7T experiment. **A. Left**: Visual stimuli used in the 7T fMRI experiment. **Right**: ROI definitions for the M and P layers of the LGN. The upper panel shows the map for the layer index. The lower panels show the ROIs for the M and P layers. **B. Upper:** The high contrast achromatic P stimulus presented to the fellow eye clearly activated the dorsal (P) layers of the LGN (Group averaged beta map). Red boxes indicate the selected FOV for slices showing in C. **Lower:** the significant T map from a representative subject (*p* < 0.05 uncorrected). **C. Left:** The left three columns show the group averaged beta maps of responses to the amblyopic eye (AE), the fellow eye (FE), and the response difference between the AE and NE conditions. The right column shows the T map of significant AE-NE voxels (*p* < 0.05 uncorrected). Maps were interpolated at 0.6mm isotropic resolution. Black dotted lines indicate the boundary of the M and P layer compartments. **Right:** Amblyopic and fellow eyes’ response in the M and P layers’ ROIs. Error bars represent standard deviation of bootstrapped distribution of the mean.

#### 1.1 3T fMRI showed selective reduction of the amblyopia eye’s response to sustained chromatic stimulus in the P layers of the LGN

As shown by Figure 1A, the chromatic P stimulus was a high spatial frequency, isoluminant red/green checkerboard pattern, reversing contrast at a low temporal frequency (0.5Hz). The M stimulus was a low contrast (30%), low spatial frequency, high temporal frequency, achromatic checkerboard flicker (10Hz). The right side of Figure 1A shows a model for the M and P layers of the LGN from Nissl stained histology (Amunts, Lepage et al. 2013) and the simulated M-P fMRI pattern (see methods for details about the simulation). Figure 1B shows the group averaged M-P beta maps of the LGN for normal, amblyopia and fellow eye groups. Different eye groups consistently showed a ventral cluster selectively biased to the M stimulus, and a dorsal cluster prefer the P stimulus. For each cluster, 30 mm^3^ voxels with the strongest response bias were shown (the highest and lowest M-P beta values for the M and P dominant clusters, respectively). This spatial pattern was consistent with the anatomy of the M and P layers of the human LGN, and consistent with our previous studies. For the amblyopia eye, response bias to the P stimulus in the P cluster was much weaker compared to the normal and fellow eye groups, suggesting a selective reduction of chromatic response in the P layers of the LGN in amblyopia.

In the ROI analysis, we extracted fMRI responses to the M and P stimuli from the M and P dominant clusters (lower panel of Figure 2B). A leave-one-subject-out procedure was used for the ROI definition to avoid the problem of “double dipping” (Kriegeskorte, Simmons et al. 2009) (see methods for details). Three-way ANOVA with within-subject factors of stimuli (M/P stimuli) and layers (M/P layers), and between-subject factor of eyes (AE/NE) showed a significant main effect of eyes (F(1,32) = 9.81, *p* = 0.0037), indicating reduced LGN response to the amblyopic eye compared to normal controls. The three-way interaction of eyes × stimuli × layers was also significant (F(1,32) = 10.82, *p* = 0.0024). Two-way ANOVAs showed significant main effect of eyes (F(1,32) = 22.27, *p* < 0.001), and interaction of eyes × stimuli in the P cluster (F(1,32) = 11.32, *p* = 0.002), but not in the M cluster (both *p* > 0.05). Post-hoc t-test showed significant AE response loss to the P and M stimulus in the P cluster (*p* < 0.001 and *p* = 0.0025, respectively), but not in the M cluster (both *p* > 0.05). These findings demonstrate that compared to normal eye’s response, amblyopia eye’s responses to both P and M stimuli were reduced in the P layers but not in the M layers, and the response loss in the P layers was more for the P stimulus. Comparison between the amblyopic eye’s and fellow eye’s responses showed similar results (main effect of eyes: F(1,16) = 13.15, *p* = 0.0023; eyes × stimuli × layers interaction: F(1,16) = 4.83, *p* = 0.043; eyes × stimuli interaction in the P cluster: F(1,16) = 8.52, *p* = 0.01; eyes × stimuli interaction in the M cluster: F(1,16) = 0.004, *p* = 0.95). Compared to the normal eye, the fellow eye’s response in the LGN showed no significant difference (main effect of eyes: F (1,32)=0.038, p=0.85, *p* > 0.05 for all interactions with the eyes). These findings clearly demonstrate a selective reduction of the amblyopia eye’s response to sustained chromatic (P) stimulus in the P layers of the LGN.

At lower magnetic field (<=3T), T2* weighted BOLD response is strongly contributed by non-specific signals from large blood vessels (Gati, Menon et al. 1997). According to the vasculature of human LGNs (Abbie 1933, Schlesinger 2012), there is a hilar region in the ventral LGN with rich vasculature. Thus the M layers’ response might be contaminated by non-specific signals from the vascular hilum. However, the dorsal P layers’ response should be less affected. And the differential maps clearly show that there still remains clear response bias (or specific signals) in the M and P dominant clusters. Another possible confounding source to the LGN’s response is the inferior pulvinar, which is closely located to the LGN and is also activated by visual stimuli. However, during data analysis, the LGN voxels were restricted within the anatomical mask of the LGN. In order to further rule out these confounding factors and to replicate our 3T fMRI findings, a 7T fMRI experiment was performed at a higher spatial resolution. At higher magnetic field, non-specific signals from draining veins in the vascular hilum should be much reduced, and specific signals from the microvessels (capillaries) will be much stronger (Gati, Menon et al. 1997, Kim 2018). With a much smaller point spread function (PSF) at 7T and at a higher spatial resolution, the influence from the inferior pulvinar should be minimal.

#### 1.2 7T fMRI revealed reduced amblyopia eye’s response to high contrast achromatic stimulus in the P layers of the LGN

According to electrophysiological studies, the role of P cells in primate vision are “double duty”: they are not only sensitive to chromatic information, but also responsive to high contrast, high spatial frequency achromatic stimulus (Derrington, Krauskopf et al. 1984, Derrington and Lennie 1984, Hubel and Livingstone 1990). Thus if there is selective activity loss of P cells in the LGN, the amblyopia eye’s response to high contrast luminance-defined stimulus should also be more reduced in the P layers than in the M layers. To test this hypothesis, and to minimize the influence from non-specific signals as in the 3T fMRI experiment, we took advantage of the resolving power of 7T fMRI to dissociate the M and P layers of the LGN according to their anatomical landmarks. Two layers of voxels were defined corresponding to the ventral and dorsal surfaces of the LGN. For the rest of voxels, we calculated a layer index as the normalized distances to the dorsal surface (ranged from 0 to 1). According the histology of human LGNs, the volume ratio between the M and P layers is about 1 to 4 (Andrews, Halpern et al. 1997). Based on this M/P volume ratio and the calculated voxel layer indices, we divided the anatomical volume of the LGN into an M and a P layers compartment (right panel of Figure 2A, and Figure S2).

The visual stimulus was a full contrast, high spatial frequency checkerboard pattern, reversing contrast at 4Hz (left of Figure 2A). According to our recent study (Qian, Zou et al. 2020), this achromatic P stimulus should strongly activate the P layers of the human LGN. Indeed, when the fellow eye was stimulated, the achromatic P stimulus strongly activated the dorsal P layers of the LGN (Figure 2B). Compared to the fellow eye, P layers’ response to the amblyopic eye was much reduced (as shown by the AE-FE beta map, left panel of Figure 2C). Significant cluster of voxels can be found in the P layer compartment (AE-NE T map, the right most column), located at the posterior part of the LGN, corresponding to the central visual field (Connolly and Van Essen 1984, Schneider, Richter et al. 2004). In the ROI analysis (right panel of Figure 2C), permutation test (see methods for details) showed significantly reduced achromatic response in the P layers (*p* = 0.026), but not in the M layers (*p* = 0.54). These findings further support selective P layers response deficits to high contrast, high spatial frequency achromatic stimulus in amblyopia.

#### 1.3 No significant difference of LGN volume between amblyopia patients and normal controls

Anatomical volumes of the LGNs were manually defined from the T1 anatomical images by two experienced researchers blind to the group labels of subjects. All subjects from the 3T and 7T experiments were enrolled in the analysis. The volumes of the left and right LGNs were averaged for each subject. The LGN volume was 132.82 ± 43.20 mm^3^ (Mean ± SD) for normal controls, and 135.26± 42.61 mm^3^ for amblyopia patients. Independent t-test showed no significant difference of LGN volumes between amblyopia patients and normal controls (t(41) = 0.205, *p* = 0.838). Also, the volume of the ipsilateral (temporal retina) and contralateral (nasal retina) LGNs to the amblyopic eye showed no significant difference (F(1, 22) = 0.111, *p* = 0.742, three bilateral amblyopia patients from the 7T data were excluded from this analysis).

### 2. Response to chromatic P stimulus was decreased to the amblyopic eye in the superficial SC, and robustly increased to the fellow eye in the deep SC

As shown by Figure 3A (upper panel), the M and P visual stimuli mainly activated the superficial SC, with smaller BOLD responses in the deeper part of the SC. It is consistent with the consensus that superficial SC mainly processes visual sensory information. The lower panel of Figure 3A show that in comparison to normal controls, the amblyopia eye’s response to the P stimulus was significantly reduced in the superficial SC, while the fellow eyes response was significantly stronger in the deeper SC (likely the intermediate layers). Compared to the fellow eye, amblyopia eye’s response was significantly weaker mainly in the superficial SC.

**Figure 3.**
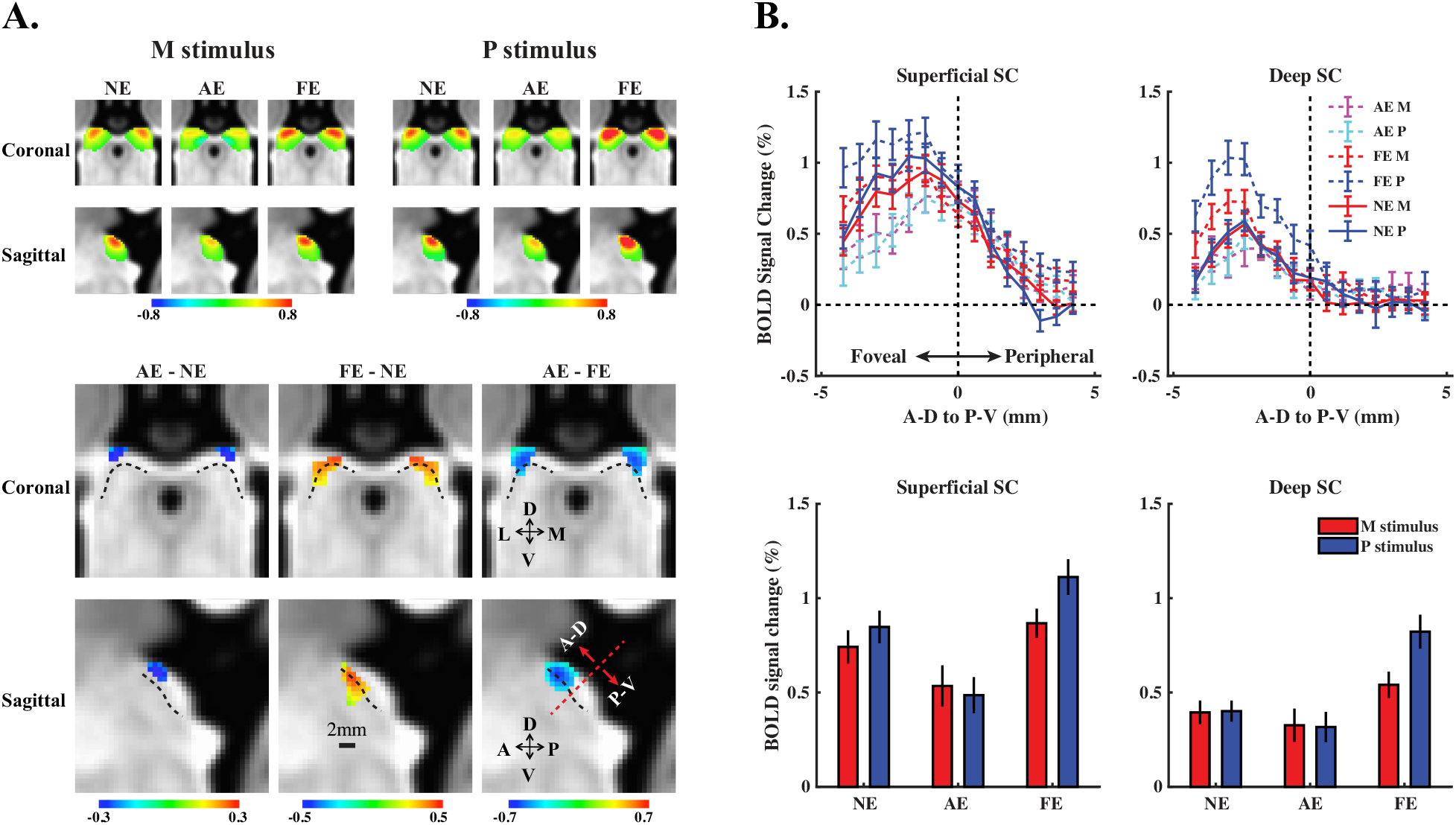
3T fMRI response to the M and P stimuli in the SC. **A. Upper:** Group averaged M and P beta maps for the normal eye (NE), amblyopia eye (AE) and fellow eye (FE) groups, respectively. **Lower:** Voxels with significantly different responses to the P stimulus for AE-NE, FE-AE, and AE-FE comparisons (uncorrected *p* < 0.05, 0.01, 0.001, respectively). Dotted lines indicate the boundary between the superficial and deep parts of the SC. **B. Upper:** BOLD response in the superficial and Deep SC from anterior-dorsal (A-D, more foveal) to posterior-ventral (P-V, more peripheral) direction. **Lower:** BOLD response in the superficial and deep SC in the anterior-dorsal half of the nucleus, corresponding to the central visual field. Error bars represent standard errors.

For ROI analysis, we first generated a normalized depth map of the SC (from 0 to 1, Figure S3), and then the SC was subdivided into a superficial and a deep part (at normalized depth = 0.5). In Figure 3B (upper panel), we plotted BOLD responses in the superficial and deep SC from anterior-dorsal to posterior-ventral direction. Results clearly show that the response differences were on the anterior-dorsal part of the nucleus, corresponding to the central visual field. Thus we calculated statistics only from the anterior-dorsal part of the SC. Three-way ANOVA of eyes (AE/NE), stimuli (M/P), and layers (superficial/deep) showed a significant interaction of eyes × layers (F(1,32) = 4.14, *p* = 0.05), and a significant interaction of eyes × stimuli × layers (F(1,32) = 4.87, *p* = 0.035). Two-way ANOVA showed that the eyes × layers interaction was significant for the P stimulus (F(1,32) = 6.7, *p* = 0.014), but not for the M stimulus (F(1,32) = 1.78, *p* = 0.19). Post-hoc t-test showed significant difference between the AE and NE response to the P stimulus in the superficial SC (t(32) = 2.2, *p* = 0.036), but not in the deep SC (t(32) = 0.69, *p* = 0.49). These results indicate that compared to normal controls, amblyopia eye’s response to the chromatic P stimulus was selectively reduced in the superficial but not in the deep SC. Three-way ANOVA of the FE and NE response showed a significant interaction of eyes × stimuli (F(1,32) = 5.28, *p* = 0.028), and significant interaction of eyes × stimuli × layers (F(1,32) = 4.63, *p* = 0.039). Two-way ANOVA showed significant main effect of eye (F(1,32) = 4.92, *p* = 0.034), and interaction of eyes × stimuli in the deep SC (F(1,32) = 14.04, *p* < 0.001), but not in the superficial SC (both *p* > 0.2). Post-hoc t-test showed significantly stronger FE response in the deep SC to the P stimulus (t(32) = xxx, *p* = 0.004), but not to the M stimulus (*p* = 0.16). These results indicate that compared to normal controls, the fellow eye’s response to the chromatic P stimulus was selectively stronger in the deep SC. Three-way ANOVA of the AE and FE response showed significant main effect of eyes (F(1,32) = 6.29, *p* = 0.023), and interaction of eyes × stimuli (F(1,32) = 13.97, *p* = 0.002), but the three-way interaction was not significant (F(1,32) = 0.035, *p* = 0.85). Thus compared to the fellow eye, the amblyopia eye’s response was selective reduced to the P stimulus in both superficial and deep SC.

### 3. Selective amblyopic response loss to chromatic P stimulus in the visual pulvinar

In Figure 4, the M and P visual stimuli significantly activated the inferior portion of the pulvinar, corresponding to the visual part of the pulvinar (Arcaro, Pinsk et al. 2015, DeSimone, Viviano et al. 2015). According to the human subcortical brain atlas from Nissl stained histology (Mai, Majtanik et al. 2015), both the ventral and the dorsal parts of the inferior pulvinar were activated. For the normal and fellow eye groups, BOLD responses in the visual pulvinar were strongly biased to the P stimulus, while the amblyopia eye’s response showed much weaker response to the P stimulus. During ROI analysis, a leave-one-subject-out procedure was used to define the ROI of visual pulvinar significantly activated by the M and P stimuli (see methods for details). The bar plot shows that the amblyopic eye’s response was strongly reduced to the P, but not to the M stimulus. Two-way ANOVA of eyes (AE/NE) and stimuli (M/P) showed significant main effect of eye (F(1,32) = 5.08, *p* = 0.031), and interaction of eyes × stimuli (F(1,32) = 13.27, *p* < 0.001). Post-hoc t-test showed significantly reduced AE response to the P stimulus (t(32) = 3.43, *p* = 0.0017), but not to the M stimulus (*p* = 0.559). Two-way ANOVA of AE and FE responses showed similar results, with significant interaction of eyes × stimuli (F(1,16) = 18.78, *p* < 0.001), and significant post-hoc t-test for the P response but not for the M response (*p* = 0.021 and 0.579, respectively). No significant difference was found between the FE and NE responses (all *p* > 0.4).

**Figure 4.**
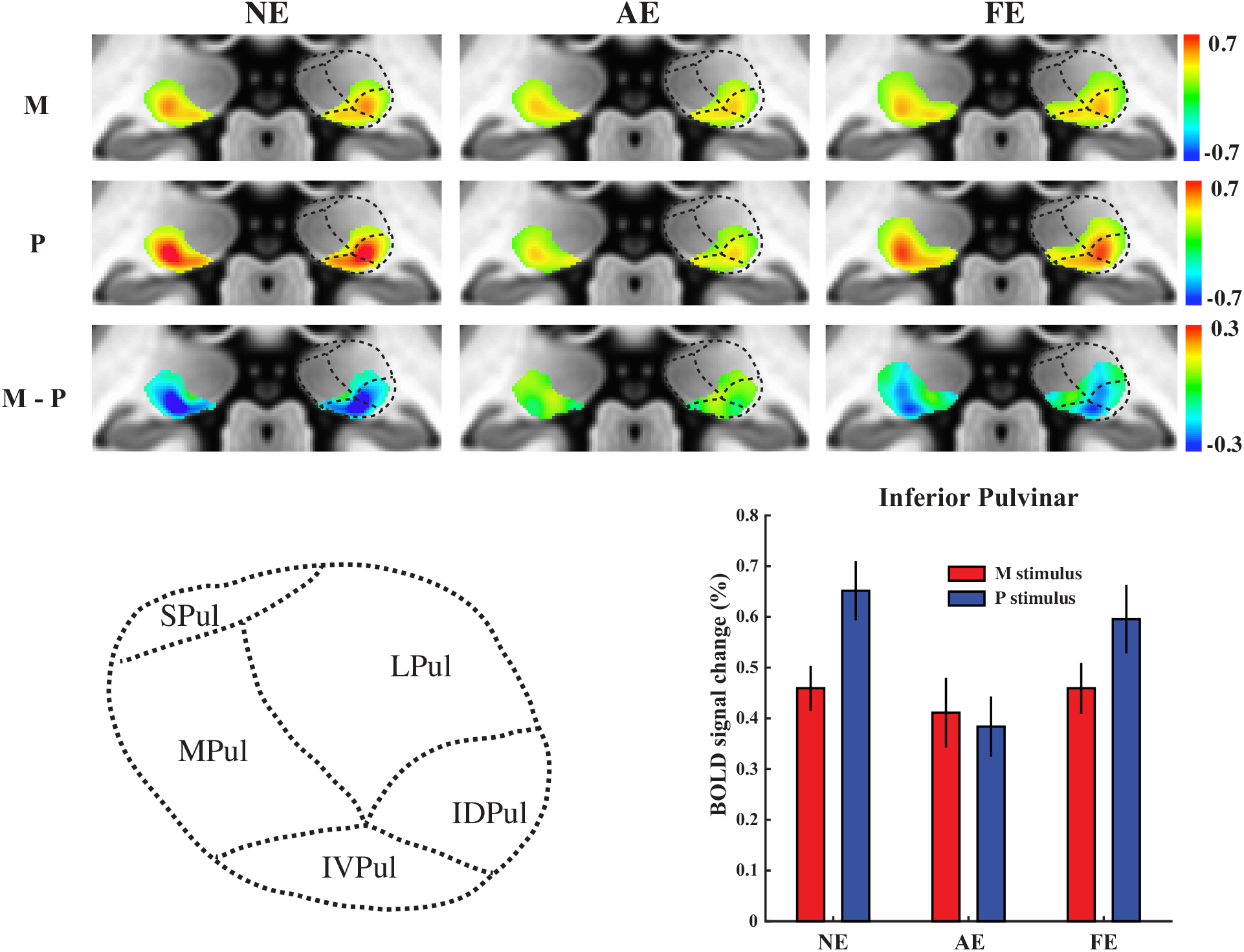
3T fMRI responses to the M and P stimuli in the visual pulvinar. **Upper:** Group averaged M-P beta maps of visual pulvinar significantly activated by the M and P stimuli (MNI y = 30mm). Maps were thresholded at M+P *p* < 0.001 uncorrected. **Lower left:** Subregions of the pulvinar according the human brain atlas based on Nissl stained histology (Mai, Majtanik et al. 2015). Black dotted lines indicate the approximate boundaries between the subregions. **Lower right:** Responses in the ROI of the inferior pulvinar. Error bars represent standard errors.

### 4. Selective amblyopic response loss to chromatic P stimulus in the early visual cortex (V1-V3) and ventral visual stream (V4/VO/LO)

Figure 5 shows the normalized fMRI responses to the amblyopic and fellow eyes (as the response ratio relative to the response of normal controls) in the subcortical and cortical visual areas. A subject-level bootstrapping procedure was used for the normalization of fMRI response (see method for details). Normalized response to the amblyopic eye was more reduced to the P stimulus than to the M stimulus in the early visual cortex (V1-V3), and in areas of the ventral visual stream (V4/VO/LO), but not in the dorsal visual stream (V3ab/MT). Notably, the amblyopic deficit of parvocellular response was much milder in the cortical areas compared to the LGN (LGNp vs. V1: p < 0.001), suggesting that the selective parvocellular functional deficit in the LGN may not be entirely inherited from cortical feedbacks. Normalized response to the fellow eye showed no significant difference with normal controls in most cortical areas. Figure S4 shows the original BOLD responses in the visual cortex to the M and P stimuli. We also examined the difference of responses from anisometropia and strabismus patient groups. A number of visual areas showed significant difference, especially in the higher order visual cortex (Figure S5). However, given the small number of subjects (10 and 7 subjects for the anisometropia and strabismus groups, respectively), and that the visual acuity was significantly different between the two groups (*p* < 0.05). These differences in BOLD response may not reliably reflect the difference of neural mechanisms between the two types of amblyopia.

**Figure 5.**
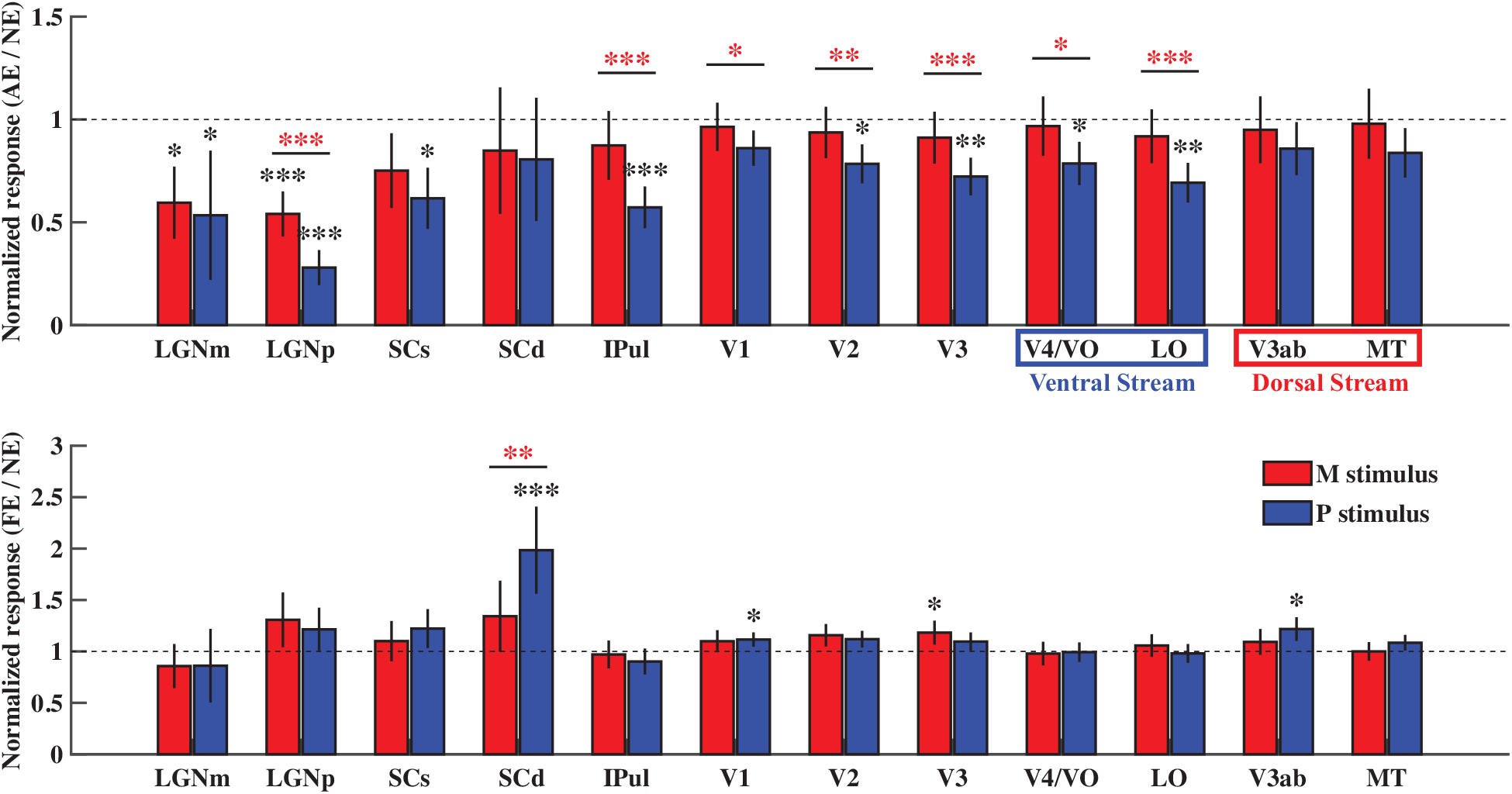
Normalized fMRI response to the amblyopic and fellow eyes in the subcortical nuclei and cortical visual areas. Upper and lower panels show the normalized response to the amblyopic eye and fellow eye, respectively (divided by the response of normal controls). Error bars indicate standard deviation of the bootstrapped distribution of the mean. Abbreviations, LGNm: M layers of the LGN, LGNp: P layers of the LGN, IPul: Inferior Pulvinar, SCs: superficial SC, SCd: deep SC. \**p* < 0.05, \*\**p* < 0.01, \*\*\**p* < 0.001.

## Discussion

In the 3T fMRI experiment, we showed that compared to the normal controls and fellow eyes, amblyopic eyes showed more reduced response to sustained chromatic stimulus than transient achromatic stimulus in the P layers, but not in the M layers of the LGN. High-resolution 7T fMRI further confirmed amblyopia eyes showed selective response loss in the P layers to high contrast, high spatial frequency luminance-defined stimulus. Robust and selective P response loss was also found in the visual pulvinar. These results provide direct in vivo evidence for selective deficits of parvocellular function in the visual thalamus of adult human amblyopia. Selective response loss to the chromatic P stimulus was also found in the early visual cortex and the ventral stream of cortical visual areas, but the effect size were much smaller compared to the LGN. Another key finding is that amblyopic eye’s response to the P stimulus was selectively reduced in the superficial SC, while the fellow eye’s response to the P stimulus was strongly increased in the deep SC. These findings clearly demonstrate functional deficits to the retinotectal pathway in amblyopia, and suggest a compensation neural mechanism in the SC to the fellow eye’s response.

In the current study, we found clear and direct evidence for selective amblyopic deficits to both chromatic and achromatic processing in the P layers of the LGN. Previous psychophysics and neuroimaging studies also suggested P-pathway selective deficits in amblyopia, but the evidence is indirect and sometimes controversial. For example, Levi and Harwerth found contrast sensitivity loss for amblyopia eye most pronounced at high spatial frequencies, while the sensitivity for low spatial and high temporal frequency stimulus was unaffected (Levi and Harwerth 1977). Using a luminance pedestal discrimination paradigm, Zele and colleagues found that spatial contrast sensitivity was similarly affected for M and P pathways (Zele, Pokorny et al. 2007, Zele, Wood et al. 2010). Contrast sensitivity for color and luminance were also found differently or similarly affected from different studies (Bradley, Dahlman et al. 1986, Davis, Sloper et al. 2006). Several neuroimaging studies found reduced suprathreshold activity for high spatial and/or low temporal frequency stimuli in the visual cortex (Barnes, Hess et al. 2001, Choi, Lee et al. 2001, Lee, Lee et al. 2001, Mizoguchi, Suzuki et al. 2005, Lerner, Hendler et al. 2006, Muckli, Kiess et al. 2006). However, spatiotemporal response properties of M and P cells are less distinctive at high contrast levels (Merigan and Maunsell 1993), and indeed some fMRI studies showed the opposite effect: more reduced cortical response to low spatial and high temporal stimuli (Hess, Li et al. 2009, Li, Yang et al. 2013). Most importantly, M and P inputs from the LGN only remain segregated in V1 input layers and merged in extra-striate cortex, thus a clear dissociation of M and P pathway-specific response from the visual cortex is not possible. A recent fMRI study found more reduced response to chromatic than achromatic stimuli in the whole LGN of amblyopia patients (Hess, Thompson et al. 2010), but the M and P layer-specific responses were not separated. Since the P cells are responsive to both chromatic and achromatic visual input, it is not clear whether there is a general functional loss of the P layers. Also given the relatively low spatial resolution of this study and a huge ROI definition for the LGN, the response could be contaminated by the inferior pulvinar which locates closely to the LGN. The inferior pulvinar indeed showed robust and selective response loss to sustained chromatic stimulus in our study. Our high resolution fMRI experiments and data analysis approach avoid the contamination from these signals.

Visual pulvinar receives driving input from the visual cortex, and regulates information transmission across cortical areas (Purushothaman, Marion et al. 2012, Zhou, Schafer et al. 2016). It also receives input from the SC and sends projections to extra-striate and posterior parietal cortex (Adams, Hof et al. 2000, Berman and Wurtz 2010, Lyon, Nassi et al. 2010). Amblyopia deficits to the function of pulvinar remain almost unknown, only one previous study showed minimal response loss of pulvinar to motion stimulus presented to amblyopic eyes (Thompson, Villeneuve et al. 2012). In the current study, we found very robust amblyopic response loss to the chromatic P stimulus in the visual pulvinar, while the response to the M stimulus was almost unaffected. This finding may suggest dysfunction of parvocellular information transmission across cortical visual areas, and/or selective P response deficit of the tecto-pulvino-cortical visual pathway.

Selective P response loss to the amblyopia eye was found in the early visual cortex (V1-V3) and ventral visual stream (V4/VO/LO), but not in the dorsal visual stream (V3a/MT). These findings further support P-pathway selective deficit in amblyopia. Compared to the LGN, cortical response deficits were much weaker, suggesting that the P response deficit in the LGN might not be entirely due to degraded cortical feedbacks. This finding was consistent with a previous study showing weaker feedforward connectivity from LGN to V1 in amblyopia (Li, Mullen et al. 2011). Thus we speculate that the amblyopic deficit might arise at the thalamic level and neural compensations may occur at the visual cortex. Another possibility is that there could be neural deficits in addition to the reduced population response in the visual cortex, such as more noisy and less stable spike trains.

To the best of our knowledge, the function of retinotectal pathway in amblyopia has never been investigated. Our results clearly showed that the amblyopic eye’s response in the superficial SC was significantly reduced to sustained chromatic stimulus. Since the superficial SC does not receive direct projection from the midget (P) ganglion cells of the retina, such chromatic response deficit should be related to feedbacks from early visual cortex (Finlay, Schiller et al. 1976, White, Boehnke et al. 2009). According to a theory of attention and recent empirical studies (Zhaoping 2008, White, Berg et al. 2017, White, Kan et al. 2017, Yan, Zhaoping et al. 2018), cortical feedbacks to the superficial SC might be related to the representation of attention saliency map. Our results may suggest a degraded attention saliency map in the SC of amblyopia patients. An interesting and surprising finding is the increased fellow eye’s response in the deep SC. Although we didn’t record eye movements in the scanner, several previous studies found dysfunctions of fixational drifts and microsaccades of the amblyopia eye but not of the fellow eye during monocular viewing conditions (Gonzalez, Wong et al. 2012, Subramanian, Jost et al. 2013, Chung, Kumar et al. 2015, Shaikh, Otero-Millan et al. 2016). However, it might be possible that the fellow eye had improved fixational eye movements (e.g. more microsaccades) specifically to stimulus with more spatial details, such as the high spatial frequency P stimulus used in the current study. Improved miniature eye movements of the fellow eye might help better sampling of fine spatial details to compensate acuity loss of the amblyopia eye (Rucci, Iovin et al. 2007, Kuang, Poletti et al. 2012). Another related explanation is a compensation mechanism of attention in the SC boosting the fellow eye’s response. In support of this hypothesis, several cortical visual areas did show slightly greater response to the fellow eye group compared to normal controls (e.g. V1 and V3ab in Figure 5). Future studies should link the psychophysics and neuroimaging data to understand the functional significance of these response changes of superior colliculus in amblyopic vision.

Currently there is no effective way to treat human adult amblyopia, an important reason might be due to the limited understanding about the neural mechanisms of the disease. The current study clearly demonstrate selective amblyopic deficit of parvocellular functions in the visual thalamus and reveal dysfunctions of the retinotectal pathway. These new findings might help to develop new tools for better treatment and prognosis of the disease, such as training paradigms specifically improving the amblyopic eye’s response in the parvocellular visual pathway.

## Materials and Methods

### Subjects

17 adult patients diagnosed with unilateral amblyopia (10 anisometropia and 7 strabismus patients, age 25.0±5.0 years, 4 females) and 17 gender matched healthy controls (age 33.0±5.6 years) were enrolled in 3T fMRI study. 9 adult patients diagnosed with anisometropia (6 unilateral and 3 bilateral patients, age 29.78±8.21 years, 6 females) were enrolled in the 7T fMRI study. The research followed the tenets of the Declaration of Helsinki, and all participants gave written informed consent in accordance with procedures and protocols approved by the Human Subjects Review Committee of the Eye and ENT Hospital of Fudan University, Shanghai, China. 7T participants also gave written informed consent approved by institutional review board of the Institute of Biophysics, Chinese Academy of Sciences.

The best-corrected visual acuity (BCVA) of each subject was assessed by experienced optometrist with the Snellen chart. Cover and alternative cover testing, duction and version testing, intraocular pressure testing, slit-lamp testing, indirect fundus examination after pupil dilation, and optometry were performed for each subject. The BCVAs of all amblyopic eyes was 20/30 or worse, and the BCVA of all fellow eyes was 20/20 or better. Both amblyopia and control participants were required to be free from a history of intraocular surgery, any eye diseases, and any systemic diseases known to affect visual function (e.g., migraine, congenital color deficiencies). None of amblyopia subjects underwent amblyopia treatment including patching treatment. All strabismic amblyopia patients had undergone strabismus surgery at least one year previously and had normal eye position and steady fixation. Eye dominance in normal subjects was determined by instructing the subject to look at a distant letter through a hole between her/his hands with their eyes alternatively closed and to report the target was visible or invisible. Dominant eye was the eye that still could maintain the letter in the hole (hole-in-card method).

### Stimuli and procedures for the 3T fMRI experiment

Visual stimuli were generated in MATLAB (Mathworks Inc.) with Psychophysics Toolbox extensions (Brainard, 1997; Pelli, 1997), running on a MacBook pro computer, and presented on a translucent screen with an MRI compatible projector at a resolution of 1024 × 768 @ 60Hz. Color lookup table of the display was calibrated to have linear luminance output.

As shown by Figure 1A, full-field (24 degrees in diameter) M and P biased visual stimuli differed in spatiotemporal frequency, contrast and chromaticity were used to selectively activate the M and P layers of the LGN. The M stimulus was a low-spatial-frequency sine wave checkerboard counter-phase flickering at 10Hz with 30% luminance contrast. The P stimulus was a high-spatial-frequency, isoluminant red/green square wave checkerboard reversing contrast at 0.5Hz. The P stimulus rotated clockwise or counterclockwise at 2.4 degrees/seconds to prevent adaptation. The subjective isoluminance of red and green was adjusted with a minimal flicker procedure, repeated four times for each subject in the scanner before the fMRI experiment. To identify the region of interest (ROI) of the LGN of visual pulvinar, hemifield checkerboard stimuli counter-phase flickering at 7.5 Hz were presented in-alternation to the left and right visual field every 16 seconds.

The fellow eye and the amblyopic eye of amblyopia patients were tested in separate sessions. Each session consisted of four runs for the M and P visual stimuli, two runs for the hemifield LGN localizer, and a T1 anatomical scan. Each fMRI run lasted 256 seconds, with two dummy scans before each run. The non-dominant eye of the normal controls was tested with the same experimental procedure. During fMRI scanning, subjects were instructed to maintain fixation and pay attention to the visual stimuli. A black eye patch covered the non-test eye, and the test eye wore glasses to correct refractive errors.

### Stimuli and procedures for the 7T fMRI experiment

Full contrast checkerboard stimuli counter-phase flickering at 4Hz was used and the diameter is 10 degree (Figure 2A). The left and right eyes of amblyopia patients were tested in the same scanning session and each session consisted of six runs: three runs for the left eye and anther three for the right eye. Each functional run lasted 276 seconds, with 16 seconds of visual stimulation (8 blocks) interleaved with 16 seconds of fixation period (9 blocks). Two dummy scans were collected in the beginning of each run and were discarded during data analysis. During the experiment, the subjects were instructed to maintain fixation and to pay attention to the stimulus. Both eyes wore glasses to correct refractive errors. A black eye patch covered the non-tested eye. After one eye finished scanning, subject switched the eye patch in the scanner by themselves.

### MRI Data acquisition

3T MRI data were collected with a 3T scanner (Siemens, Verio) at the Eye and ENT Hospital of Fudan University using a twelve-element head matrix coil. A reduced field of view (FOV) gradient echo planar imaging sequence was used to acquire functional images (2mm isotropic voxels, 26 axial slices, 64×64 matrix with 2mm in-plane resolution, 2mm thickness, TR/TE = 2000/28 ms, flip angle = 90°, a saturation band was placed on the frontal regions to avoid image wrap-around). Reducing the FOV significantly reduces the echo train length of EPI, thus alleviates image distortion and blur of high resolution imaging (Heidemann, Anwander et al. 2012). Anatomical volume was obtained with a T1 MPRAGE sequence (1mm isotropic voxels, 192 sagittal 1mm-thick slices, 256 × 256 matrix with 1mm in-plane resolution, TR/TE = 2600/3.02 ms, flip angle = 8°).

7T fMRI data were acquired on a 7T MRI scanner (Siemens Magnetom) with a 32-channel receive 1-channel transmit head coil (Nova Medical), from Beijing MRI center for Brain Research (BMCBR). Gradient coil has a maximum amplitude of 70mT/m, 200us minimum gradient rise time, and 200T/m/s maximum slew rate. Functional data were collected with a T2*-weighted 2D GE-EPI sequence (26 axial slices, TR = 2000 ms, TE = 22 ms, image matrix =150×150, FOV = 180×180 mm, GRAPPA acceleration factor = 2, Flip angle=80°, partial Fourier = 6/8, phase encoding direction from A to P). Before each functional scan, five EPI images with reversed phase encoding direction (P to A) were also acquired for EPI distortion correction. Anatomical volumes were acquired with a T1-MP2RAGE sequences at 0.7 mm isotropic resolution (256 sagittal slices, centric phase encoding, acquisition matrix=320×320, Field of view = 224×224 mm, GRAPPA=3, TR = 4000 ms, TE=3.05 ms, TI1 = 750ms, flip angle = 4°, TI2 = 2500 ms, flip angle = 5°). Subjects used bite bars to restrict head motion.

### MRI Data analysis

#### Preprocessing

Preprocessing of functional data was done in AFNI (Cox 1996), including the following steps: physiological noise removal with retrospective image correction, slice timing correction, EPI image distortion correction with nonlinear warping (PE blip-up, for 7T data only), rigid body motion correction, alignment of corrected EPI images to T1-weighted anatomical volume (cost function: lpc), and per run scaling as percent signal change. To minimize image blur, all spatial transformations were combined together and applied to the functional images in one resampling step. General linear models with a fixed HRF (Block4 in AFNI) were used to estimate BOLD signal change from baseline for each stimulus condition. Motion parameters were included as regressors of no interest. Subcortical data were transformed to MNI space using advanced normalization tools (ANTs) (Avants, Tustison et al. 2011). We used the symmetric MNI template from CIT168 atlas for nonlinear warping (Pauli, Nili et al. 2018). Functional data of the subcortex were resampled to 0.6mm isotropic in MNI space. MNI space data were used for the ROI analysis of the SC and Pulvinar, native space data (up-sampled to 1mm, and 0.6mm isotropic for the 3T and 7T data, respectively) were used for the LGN ROI analysis.

#### ROI definition and analysis

##### LGN

For the 3T experiment, ROIs of the LGN were defined as voxels significantly activated by the hemifield localizer (uncorrected *p* < 0.05) within the anatomical mask of the LGN. Anatomical ROIs of the LGN were defined by two experienced researchers from T1-MPRAGE images, in which the LGN appeared darker than surrounding tissues. Care was taken not to include activations from the inferior pulvinar. Beta values of the M and P stimuli were extracted from all voxels of the ROI and entered group-level analysis in MATLAB. For each eye group, the coordinates of the left LGNs were mirror-flipped to the right, and all LGNs were registered to the center of mass location. A median LGN mask was then generated (voxel has values from half of the LGNs) as the registration template. Then a twelve-parameters linear (affine) transformation to the median LGN template was performed on each LGN. This procedure ensured approximate matching of orientation and size of the LGNs. Then group level analysis was performed on this “standard space”. M and P ROIs were defined from the M-P beta map. In order to avoid the problem of ‘double dipping’, we used a leave-one-subject-out procedure to define the M and P layer ROIs from the M-P beta map. For each subject, the M-P beta map was generated from the data of the rest subjects. The M ROI was identified as 30 voxels with strongest response bias to the M stimulus (highest M-P values), while the P ROI was defined as 30 voxels with strongest response bias to the P stimulus (lowest M-P values). ROI of the LGNs for the 7T experiment were defined only from the T1-weighted anatomical volumes. The ROI registration procedure was similar to the 3T experiment, except that an LGN template from the MNI space was used. A normalized layer index map was generated on the LGN template. Two layers of voxels were defined corresponding to the ventral and dorsal surfaces of the LGN. For the rest of voxels, we calculated a layer index as the normalized distances to the dorsal surface (ranged from 0 to 1). The ROIs for the M and P layers were determined from the layer index map according the volume ratio of M/P layers of human LGNs (M/P = 1/4) (Andrews, Halpern et al. 1997).

##### SC

ROI analysis for the SC was performed with MNI space data from the 3T experiment. ROIs of the SC were defined manually from the CIT168 MNI T1 template. To generate the depth map of the SC, two layers of voxels were first defined from the superficial and deep surface of the SC. Then a normalized depth map was calculated for each voxel as the ratio of shortest distances to the superficial and deep surfaces (from 0/superficial to 1/deep). The volume of the SC was split into a superficial and a deeper part at depth = 0.5. To generate the AD to PV slice profile in Figure 3B, the peak response in the superficial and deep ROIs were selected for each slice.

##### Pulvinar

ROI analysis for the pulvinar was also performed with the MNI space data from 3T. Anatomical ROIs of the Pulvinar were defined from the CIT168 MNI template. Spatial smoothing was performed within the ROI with a 1.2mm FWHM Gaussian filter. For each subject, a leave-one-subject-out-procedure was used to define the functional ROI, 300mm^3^ voxels with the strongest M+P beta values from the rest subjects in the group were selected as the ROI.

##### Visual cortex

Retinotopic ROIs of cortical visual areas were defined on the cortical surface according to the polar angle atlas from the 7T retinotopic dataset of Human Connectome Project (Benson, Jamison et al. 2018).

##### Statistics

A subject-level bootstrapping procedure was used for calculating the statistics of 7T response in Figure 2C (right), and the normalized fMRI response in Figure 5. For each permutation, the data for a group of subjects were resampled with replacement. Then the group averaged fMRI responses for different eye groups were calculated. The bootstrapping procedure was repeated 10^6^ times. A null distribution was generated by shifting the mean of bootstrapped distribution to the test value, and a two-sided p value was derived from the null distribution.

## Acknowledgements

This study was funded by Beijing Science and Technology Q2 Project (grant nos. Z171100000117003 and Z181100001518002); Ministry of Science and Technology of China (grant no. 2019YFA0707103); Bureau of International Cooperation, Chinese Academy of Sciences (grant no. 153311KYSB20160030) and National Natural Science Foundation of China (grant no. 31871107, 31930053 and 81500752).

## Supplemental Materials

### M-P pattern simulation for the 3T BOLD fMRI experiment

As shown by Figure S1, we first manually identified each individual layer of the LGN from Nissl stained brain slices at 20μm resolution (Amunts, Lepage et al. 2013). The neural image for each stimulus was generated by labeling in the corresponding layers with a normalized response magnitude: 1 for the P layers, and 4 for the M layers, based on much higher sensitivity of the M cells (Wiesel and Hubel 1966, Derrington and Lennie 1984). Then spatial smoothing with a 3.5mm FWHM gaussian filter was applied on the neural pattern to simulate the blurring effect of GE BOLD point spread function at 3T (Engel, Glover et al. 1997). Then the BOLD image was resampled to the resolution of fMRI measurement (2mm isotropic) and up-sampled to 0.6mm isotropic with cubic interpolation as in the fMRI data analysis, finally a differential image was generated by subtracting the patterns between the M and P stimulus conditions.

**Figure S1.**
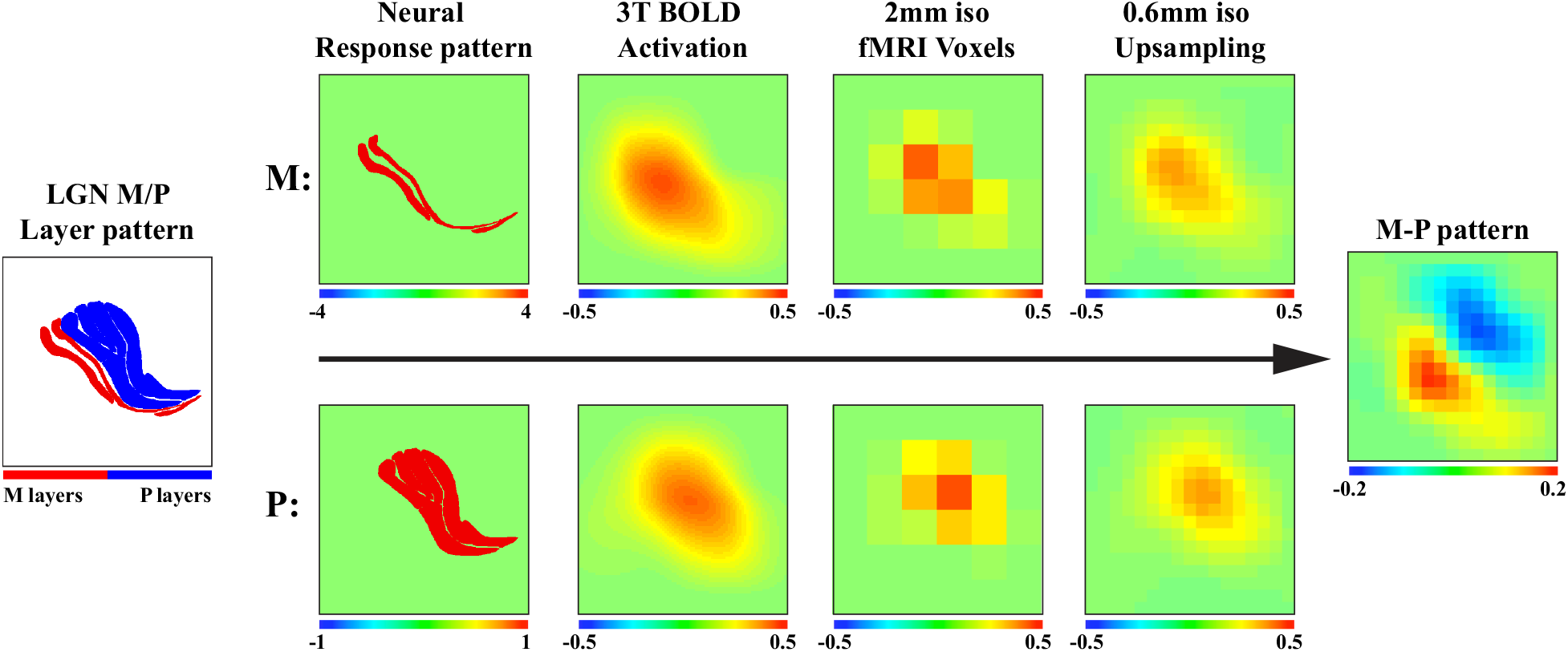
M-P pattern simulation for the 3T BOLD fMRI experiment.

**Figure S2.**
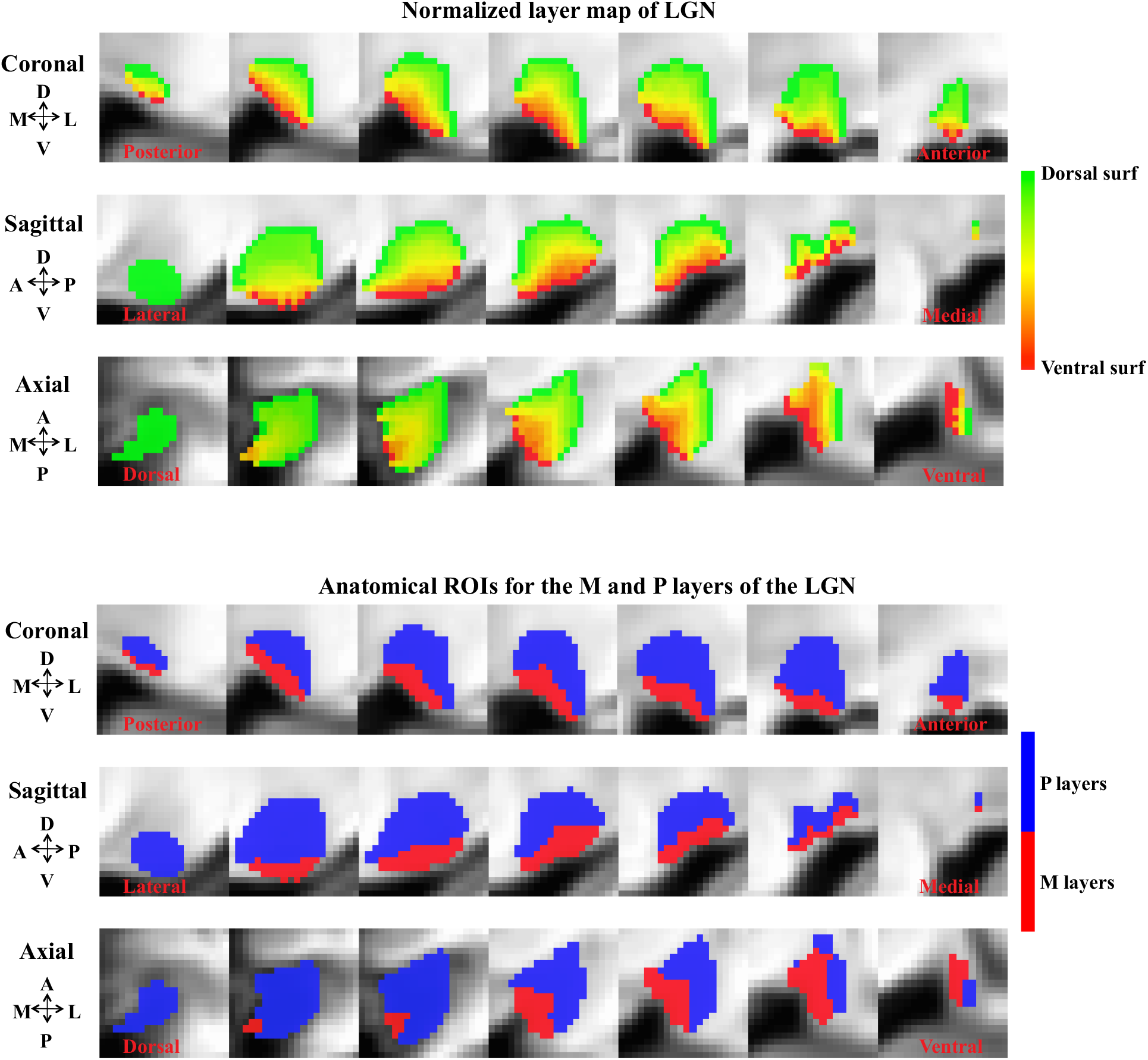
Normalized layer index map and ROIs for the M and P layers of the LGN (for the 7T experiment). Two layers of voxels were defined corresponding to the ventral and dorsal surfaces of the LGN. For the rest of voxels, we calculated a layer index as the normalized distances to the dorsal surface (ranged from 0 to 1). The ROIs for the M and P layers were determined from the layer index map according the volume ratio of M/P layers of human LGNs (1:4)

**Figure S3.**
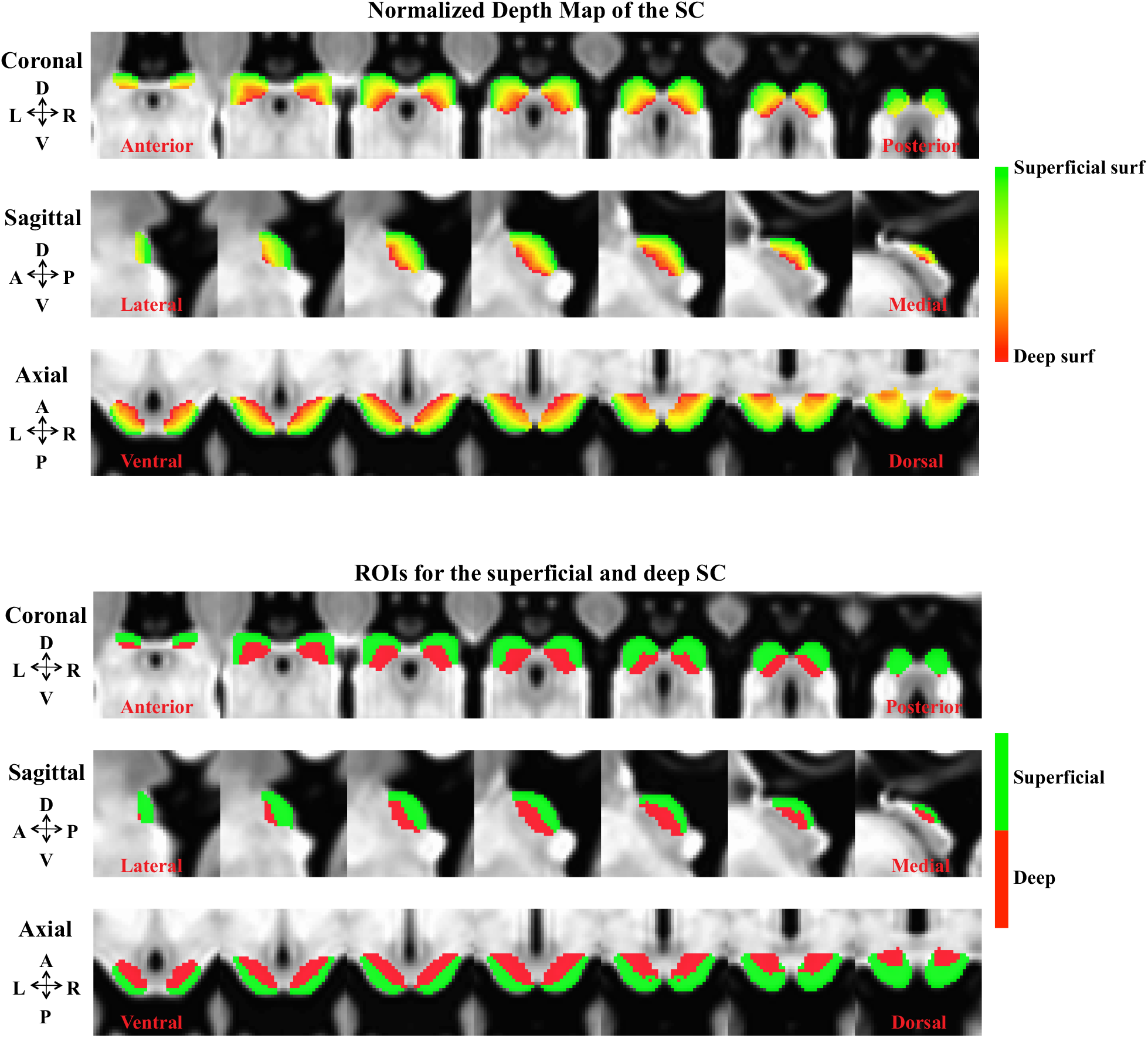
Normalized depth map and the superficial and deeper layers ROIs for the SC. Two layers of voxels were first defined from the superficial and deep surface of the SC. Then a normalized depth map was calculated for each voxel as the ratio of shortest distances to the superficial and deep surfaces (from 0/superficial to 1/deep). The volume of the SC was split into a superficial and a deeper part at depth = 0.5.

**Figure S4.**
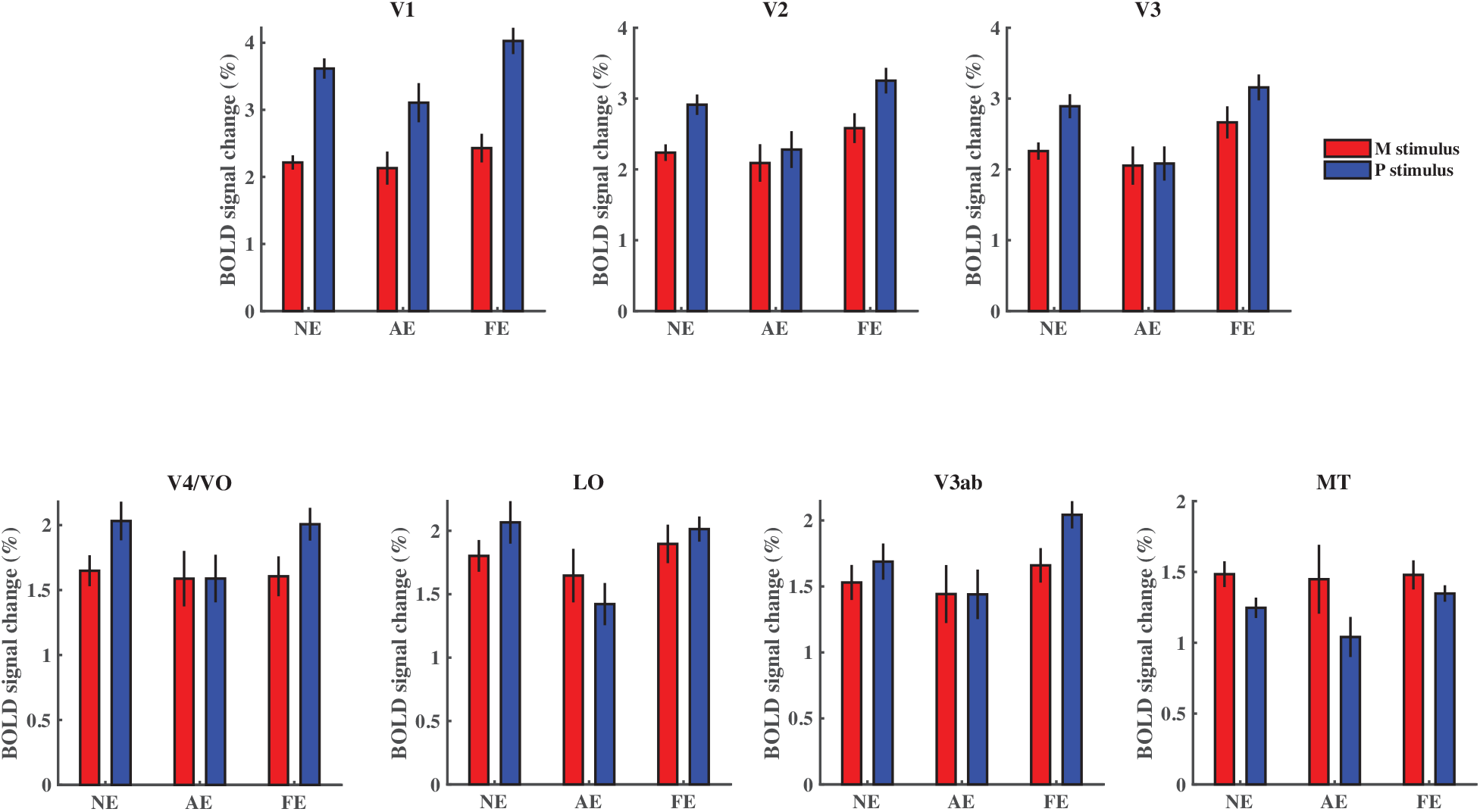
3T fMRI response to the M and P stimuli in cortical visual areas. Significant two-way interaction between eyes (AE/NE) and stimuli (M/P) was found in V1, V2, V3, V4/VO, and LO (all *p* < 0.05), but not in V3ab and MT (both *p >* 0.2).

**Figure S5.**
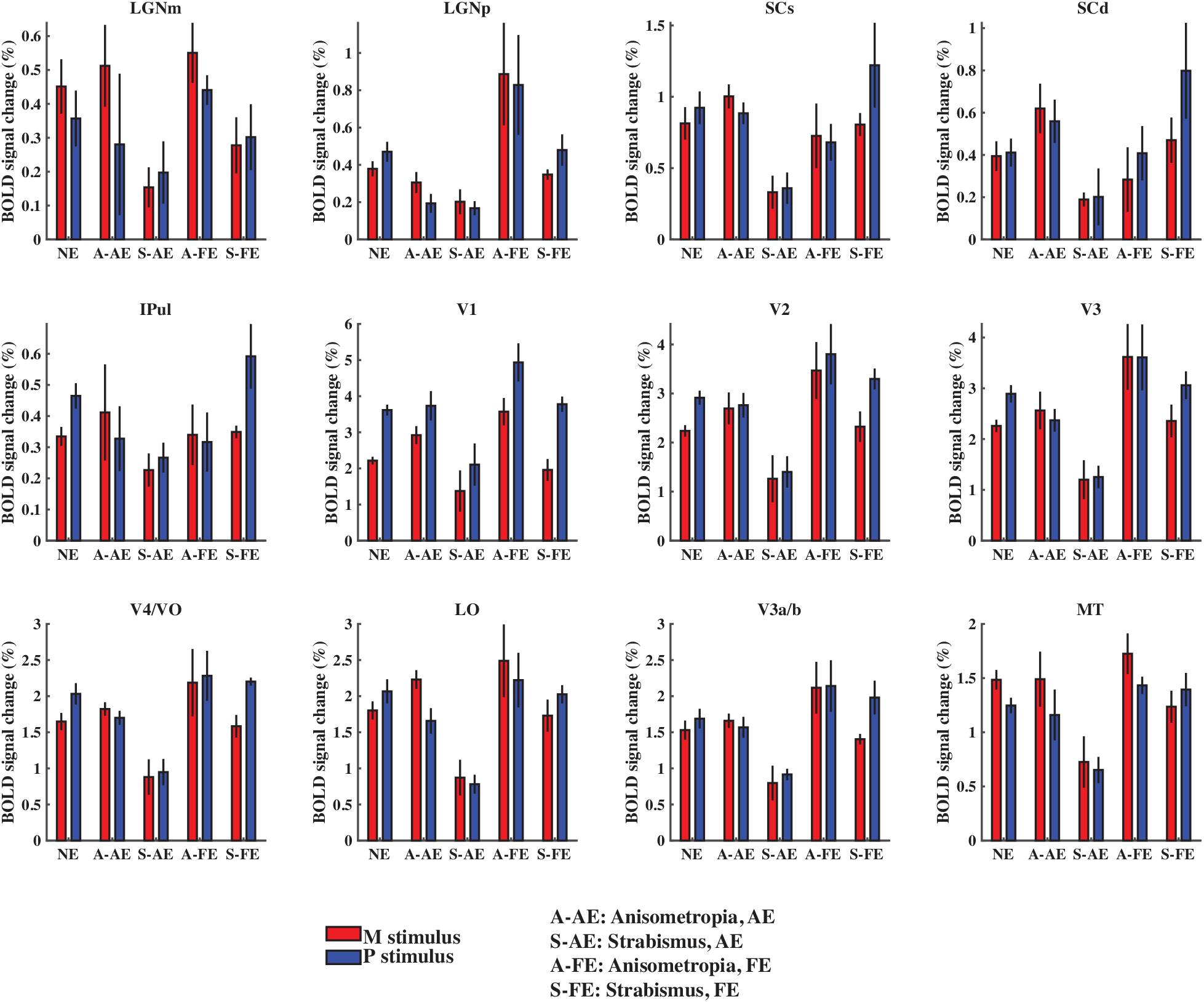
Subcortical and cortical responses for anisometropia and strabismus patients. Significant difference between the response to the amblyopic eye of Anisometropia and Strabismus patients can be found in higher order visual areas: V3, V4/VO, LO, and MT (*p* < 0.05 for interaction of groups (anisometropia/Strabismus) and eyes (AE/FE)).

